# Assessing the Incidence and Severity of Field Diseases in Marigold (*Tagetes erecta*) Cultivation: Results from Jashore District, Bangladesh

**DOI:** 10.1101/2024.10.25.620302

**Authors:** Anannya Shome, Abu Noman Faruq Ahmmed, Rabita Zaman

## Abstract

A research investigation was carried out in the Jashore district of Bangladesh to study diseases affecting marigold plants, while the laboratory analyses were conducted in the Plant Disease Study Facility, Sher-e-Bangla Agricultural University, Dhaka. The experiment was conducted by surveying three farmers from each of the sixteen villages across four unions in Jhikargacha Upazila to document the occurrence and severity of diseases under natural field conditions. The fungal pathogens surveyed in these diseases were six: leaf spot, botrytis blight, flower bud rot, stem rot, foliage blight, white mold-soil-borne diseases. The most important microflora reported to be associated with these diseases include *Alternaria alternata, Botrytis cinerea, Alternaria dianthi, Fusarium oxysporum, Curvularia lunata*, and *Sclerotinia sclerotiorum*. Of these, major diseases included leaf spot, botrytis blight, foliage blight or flower and leaf blight, and flower bud rot, while minor diseases included stem rot and white mold. Disease incidence ranged from 0-54.33%, while disease severity varied from 1-24%. The paper is the first record of marigold pathogens in Bangladesh and will be highly useful during disease identification, and it would lead to the better cultivation of marigolds in this area.

## 1. Introduction

Flowers are associated with beauty and purity. The demand for flowers is increasing rapidly due to their aesthetic appeal. They are in use on weddings, birthdays, religious offerings, and in social events; they play a crucial role in the development of various industries and economies [1]. Today, flower cultivation is a lucrative business that yields a higher return than many other crops. The global flower market is unfolding, and Bangladesh’s favorable climate for the growth of flowers has made the flower sector an emerging one that is contributing much to Bangladesh’s GDP growth and job creation [2]. Bangladesh can avail vast opportunities in the international demand for flowers, as the country is one of more than 90 countries participating in the international floriculture trade [3]. Flowers hold great significance in Bangladesh, promising good returns and engaging thousands of farmers, particularly in the Jashore district alone where 9,000 hectares are covered [4,5]. The domestic market for flowers is estimated to be Tk. 300 crore, whereas the international sales reportedly reached Tk. 16,000 crore [6]. Despite facing challenges within the flower sector, like infestation by pests, growing flowers remains lucrative-257% average annual profit margin in marigold flowers within a six-month harvest cycle [7,8]. The Flower Society of Godkhali, Jashore, estimated that a quantity of around USD 54 crores is produced every year in Godkhali, Jeshore alone and stands at USD 100 crores in total flower business amount. Gross margins of flower were Tk.1,359,824.20 per hectare [9].

Marigold belongs to the family Asteraceae. It is one of the commercially promising flowers because it is easy to cultivate and has adaptation and increased demand in the Indian subcontinent [10]. The marigold is among the most popular ornamental flowers of Bangladesh. It is commercially grown in Jhikargacha, Jhenaidah, Gazipur, and Chittagong areas of Bangladesh. This is because there is a high demand for it due to its ease of cultivation, being marketable as loose flowers, as well as being used in landscaping. Both the leaves and flowers of marigold plant have phenolic activities, as well as antioxidant activities, due to which they are used for medicinal purposes [11].

Generally, diseases are considered among the major constraints for the growing of marigold, seriously causing crop losses. Plants are said to reduce yields globally by about 13% because of the presence of a disease [12]. Other constraints include a lack of knowledge about improved production methods, expensive hybrid seed costs, seed quality issues, and insect infestations, in addition to other severe biotic and abiotic stresses that finally affect marigold farming [13].

The present study involves the identification and assessment of marigold diseases in the Jashore district of Bangladesh. The pathogens of six major diseases, namely leaf spot, botrytis blight, flower bud rot, stem rot, foliage blight and white mold of marigold plants, were identified by cultural, morphological, and microscopic methods in the Plant Pathology Laboratory of Sher-e-Bangla Agricultural University. It also measures the incidence and severity of these diseases under natural field conditions, thus pointing out the variability in the prevalence of diseased conditions. This review work provides the first comprehensive study to assess pathogens of marigold in Bangladesh.

## 2. Materials and methods

### 2.1. Experimental site

The field experiments were conducted in the Jashore district of Bangladesh, where the fieldwork was done at 16 locations of 4 unions of Jhikargacha Upazila. The laboratory work and seed health studies were carried out at the Plant Disease Clinic, Sher-e-Bangla Agricultural University, Dhaka. The site is at 89°08′E and 23°06′N, within the High Ganges River Floodplain (AEZ NO. 11) and lies at 9 meters above sea level. Intensive surveys were undertaken at these places to record the data on diseases of marigold.

### 2.2 Experimental Period

The experiment was carried out during the period from February to May 2019. Testing for seed health was conducted in February to April of 2019.

### 2.3 Weather conditions

The fieldwork was carried out in winter, during February 2019 and the recorded average climatic condition stood at a temperature of 22°C with no rainfall and 45% humidity.

### 2.4 Soli conditions

The general soil type was moderately organic, with a medium fertility status, the tested fields being above the flood level and having adequate sunlight throughout the experiment period.

### 2.5 Data collection

The field research was conducted in the natural environment, directly on the plant surfaces. The data was collected using a survey form on the diseases of plants, which included their symptoms and those affected parts of the plant. Disease distribution, status, percentage of occurrence, and intensity were other data collected. Data on each field pertaining to disease occurrence and intensity was gathered by collecting thirty plants sampled from five locations. The values on the occurrence of disease and its intensity were computed in the manner presented by formula [14]

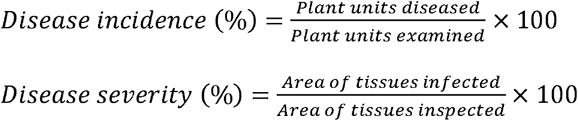

### 2.6 Isolation and identification of the responsible organisms

Diseases portions of the plants showing distinct symptoms were collected from different marigold plants in the field. Collected samples were temporarily stored in sealed poly bags with labels and brought to the Plant Disease Study laboratory. They were stored at 4°C until further analysis. The collected specimens were studied, and the causal microorganisms were isolated. Planting of the tissues was done by washing small fragments of the infected plant sections properly with sterile water and surface-sterilized with 1% Mercuric Chloride (HgCl2). The surface-sterilized fragments were cultured on the medium PDA, incubated at 25±2ºC for 6-7 days, and hyphal tips from the growing colonies were transferred on fresh PDA media for pure cultures. Growth observed under the microscope showed morphological features.

Several steps identified the causal organisms. A field survey was conducted to visually assess the symptoms and monitor its progress as well as observe the signs of pathogens under environmental conditions [15,16]. Then, diseased tissue samples were directly observed under stereoscopic and compound microscopes and fungal structures were analyzed for identification. Fungi from infected tissues were cultured onto the moist incubation technique by incubating sterilized and diseased plant parts on blotting paper for seven days. Thereafter, fungal growth was studied under a stereomicroscope. Structures were mounted on slides for detailed study under a compound microscope. Identification of pathogens was done using literature. In the final phase of culture, PDA medium was inoculated with causal organisms and fungal growth and sporulation were observed after incubation at room temperature for three days. Single spore or hyphal tip techniques were employed to obtain pure cultures and slides were prepared for microscopical analysis. Identification of pathogens was done with the help of literature and CMI descriptions [17-22].

### 2.7 Statistical analysis

The data were subjected to descriptive statistics analysis using the software STATISTIX-10, with a one-way variance analysis applied. Every village was considered as a separate treatment, and three fields per village as replications. Statistical significance level used is at P ≤ 0.05, and means were compared by LSD test. Data on disease incidence and severity were square root transformed to meet the assumptions of ANOVA. Missing data points were omitted to ensure the accuracy of the recorded data, whereas outliers were detected, assessed, and, when necessary, withheld to be representative of the natural variation measured in the field. The survey data also underwent an average mean test.

## 3. Results

### Study of Symptom Analysis

Among them, six diseases of marigold plants were identified by visual scanning during the field survey that was authenticated by symptomological studies. Such diseases were caused by particular fungal pathogens. Symptoms of the identified diseases are presented in Table 1 and the visual presentation of symptoms is shown in Fig. 1.

**Table 1.** Field observations of marigold diseases in Jhikargacha Upazila, Jashore district, Bangladesh.

| Sl. | Name of disease | Observable symptom | Disease type |
| --- | --- | --- | --- |
| 1. | **Leaf Spot** | Marigold plants with leaf spot disease developed smoky gray to black spots that grew into dark brown areas with purple edges. The infection started on lower leaves, spread to the upper ones, and caused wilting and drying. In severe cases, it led to plant death, and in moist conditions, dark mold formed on the affected areas. | **Fungal** |
| 2. | **Botrytis blight** | Botrytis blight, or "gray mold," is marked by gray, fuzzy mold on flowers and leaves. It causes leaf and flower spots, stem lesions, dieback, and stem rot, leading to plant collapse. Affected buds often fail to open, and open flowers decay and drop prematurely. | **Fungal** |
| 3. | **Flower Bud Rot** | Infected young flower buds shriveled, turned deep brown, and dried up. While mature buds showed milder symptoms, they also failed to open due to the pathogen's impact. The pathogen affected the leaves as well, causing blight characterized by brown necrotic spots on the margins and tips of older leaves. | **Fungal** |
| 4. | **Stem Rot** | Initially, brown spots formed on the stem, followed by necrosis. Older leaves yellowed and eventually died. If the infection occurred after flowering, flowers showed abnormalities in size, shape, and color. As the disease progressed, the stem broke down due to necrosis, and flower stalks curved into an S-shape. Common symptoms included firm brown to black stem rot, yellowing, browning, and premature death of leaves, along with stem decay. | **Fungal** |
| 5. | **Foliage Blight** | Foliage blight in marigolds caused outer petals to blacken and shrivel, sometimes starting from the center. Injured petals worsened the blight, and severe cases affected all leaves with small brown spots appearing. | **Fungal** |
| 6. | **Foot and Root Rot** | Foot and root rot in marigolds caused stunted growth, off-color foliage, and white fungal colonies on leaves, flowers, and stems. Plants showed yellowing, wilting, and eventually died, with root and crown decay being common after flowering. | **Fungal** |

**Table 2.** Prevalence and severity of different marigold diseases in Jhikargacha Upazila, Jashore district, Bangladesh.

| Location<br>(Villages) | Leaf Spot |  | Botrytis blight |  | Flower Bud Rot |  | Stem Rot |  | Foliage Blight |  | Foot and Root Rot |  |
| --- | --- | --- | --- | --- | --- | --- | --- | --- | --- | --- | --- | --- |
|  | DI(%) | DS(%) | DI(%) | DS(%) | DI(%) | DS(%) | DI(%) | DS(%) | DI(%) | DS(%) | DI(%) | DS(%) |
| Godkhali | 31.33 g | 14.00 ab | 34.00 a-d | 21.00 a | 23.00 b-e | 9.33 a-c | 26.00 a | 10.33 ab | 42.00 a | 17.00 a | 18.67 a-c | 7.33 a-c |
| Patuapara | 54.33 a | 14.00 ab | 37.67 a | 15.00 a-c | 34.67 ab | 16.00 a-c | 16.67 a-d | 6.67 a-c | 41.00 a | 16.00 a | 7.67 bc | 3.00 bc |
| Sadirali | 38.33 c-g | 13.00 ab | 33.33 a-d | 12.67 b-d | 22.33 b-e | 6.03 bc | 18.67 a-d | 11.67 a | 41.67 a | 17.33 a | 25.67 a-c | 9.00 a-c |
| Belemath | 38.33 c-g | 10.33 b | 26.00 b-e | 9.33 b-d | 27.67 a-e | 17.33 ab | 21.67 a-c | 8.67 a-c | 30.00 ab | 12.00 ab | 20.67 a-c | 6.00 a-c |
| Dhalipara | 41.67 b-f | 14.00 ab | 29.00 a-e | 8.67 cd | 26.67 a-e | 13.33 a-c | 26.33 a | 8.70 a-c | 19.67 bc | 8.33 ab | 15.00 a-c | 3.33 bc |
| Panisara | 34.00 e-g | 12.00 ab | 29.67 a-e | 13.67 a-d | 25.33 a-e | 9.33 a-c | 20.00 a-d | 10.00 ab | 32.33 ab | 14.00 a | 24.67 a-c | 7.00 a-c |
| Sayedpara | 33.33 fg | 11.33 ab | 38.33 a | 13.70 a-d | 15.00 de | 9.67 a-c | 10.67 b-d | 2.67 c | 24.67 a-c | 8.67 ab | 5.00 c | 1.67 c |
| Nilkonthonogor | 43.67 b-d | 10.32 b | 27.00 b-e | 9.67 b-d | 16.00 de | 11.33 a-c | 9.33 cd | 3.34 bc | 34.33 ab | 9.33 ab | 12.67 a-c | 4.67 a-c |
| Kuliya | 33.00 fg | 14.00 ab | 15.33 f | 6.67 d | 30.67 a-d | 14.67 bc | 15.67 a-d | 7.00 a-c | 32.33 ab | 15.00 a | 21.00 a-c | 7.10 a-c |
| Gaburapur | 35.00 d-g | 9.33 b | 24.67 c-e | 9.67 b-d | 27.00 a-e | 11.33 a-c | 18.00 a-d | 9.33 a-c | 5.67 c | 1.00 b | 14.00 a-c | 6.10 a-c |
| Hariya | 30.00 g | 11.67 ab | 28.67 a-e | 12.67 b-d | 12.00 e | 6.67 a-c | 23.67 ab | 11.67 a | 24.00 a-c | 7.67 ab | 6.33 bc | 2.33 bc |
| Nimtola | 47.67 a-c | 12.67 ab | 35.00 a-c | 10.00 b-d | 19.67 b-e | 7.67 a-c | 18.33 a-d | 8.00 a-c | 29.67 ab | 11.33 ab | 27.67 ab | 7.33 a-c |
| Chandpur | 37.33 d-g | 11.00 ab | 20.00 ef | 6.70 d | 27.00 a-e | 10.67 a-c | 25.00 a | 11.67 a | 29.70 ab | 12.00 ab | 29.67 a | 13.00 a |
| Mathuapara | 49.00 ab | 12.33 ab | 24.00 d-f | 8.67 cd | 40.66 a | 18.00 a | 22.67 a-c | 6.67 a-c | 23.33 a-c | 7.70 ab | 22.33 a | 4.00 a |
| Nirbashkhola | 34.33 d-g | 12.67 ab | 20.00 a-e | 11.00 b-d | 30.33 a-d | 10.00 a-c | 29.00 a | 13.00 a | 35.67 ab | 18.00 a | 18.33 a | 2.83 a |
| Shiorda | 43.44 b-e | 18.00 a | 35.33 ab | 17.00 ab | 31.33 a-c | 14.67 a-c | 7.00 d | 3.33 bc | 30.00 ab | 11.00 ab | 13.00 a | 2.67 a |
| LSD 0.05 | 9.48 | 6.84 | 10.52 | 8.00 | 16.11 | 11.94 | 13.60 | 7.17 | 19.21 | 12.10 | 22.11 | 3.50 |
| CV (%) | 14.60 | 32.79 | 21.66 | 41.45 | 37.86 | 64.03 | 42.40 | 51.96 | 38.41 | 62.45 | 68.93 | 66.83 |

\*DI- Disease incidence, DS- Disease severity, Means followed by the same letters in a column do not differ at 5% level of significance by LSD

**Figure 1.**
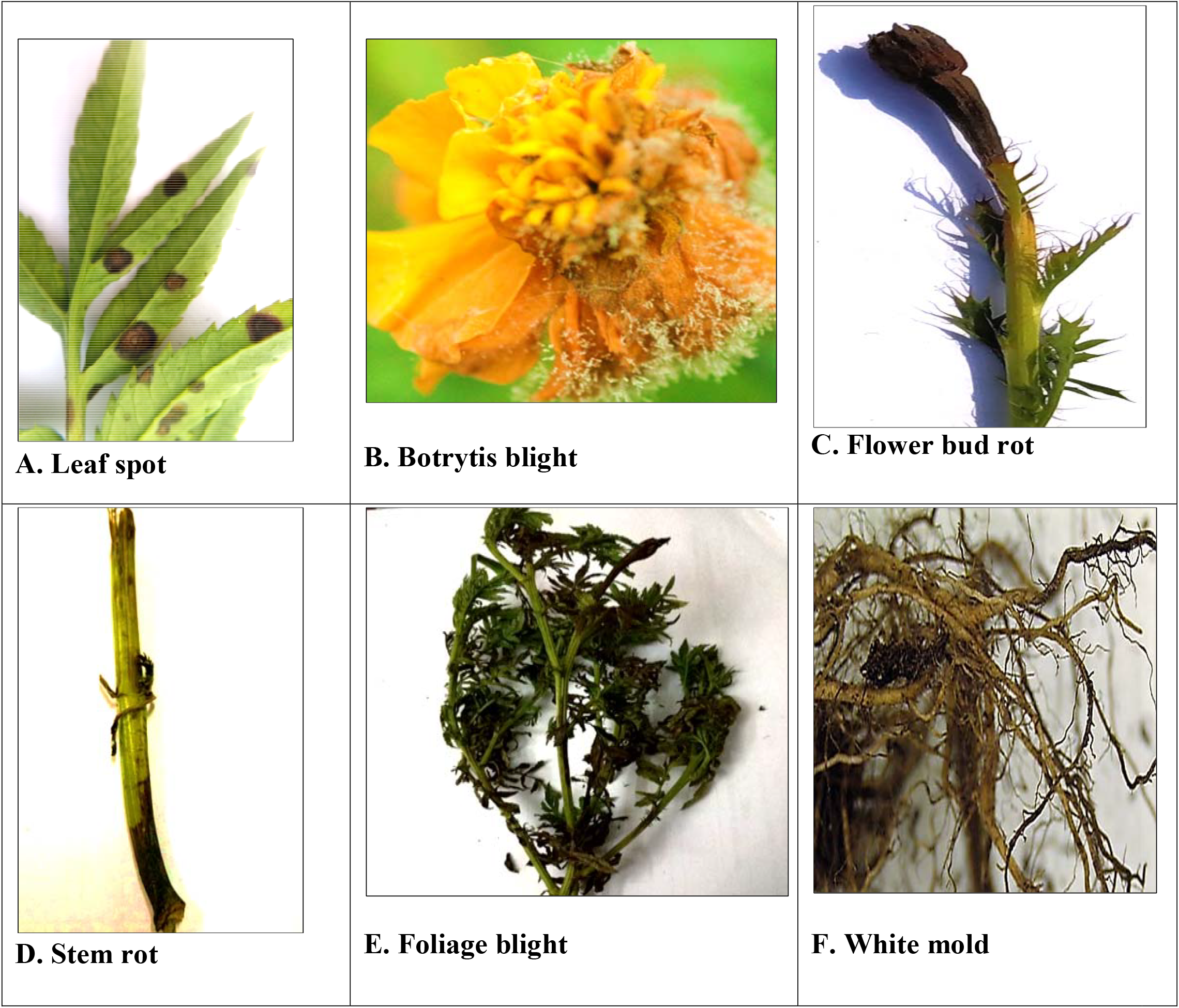
Symptoms observed in the field for different marigold diseases **A**. Leaf spot, **B**. Botrytis blight, **C**. Flower bud rot, **D**. Stem rot, **E**. Foliage blight, **F**. White mold

### Identifying the causal organism

Laboratory studies on various diseased plant parts, isolated, resulted in the identification of six pathogens by observing the pure cultures of the causal organisms, whose microscopic characteristics are shown in Fig. 2.

**Figure 2.**
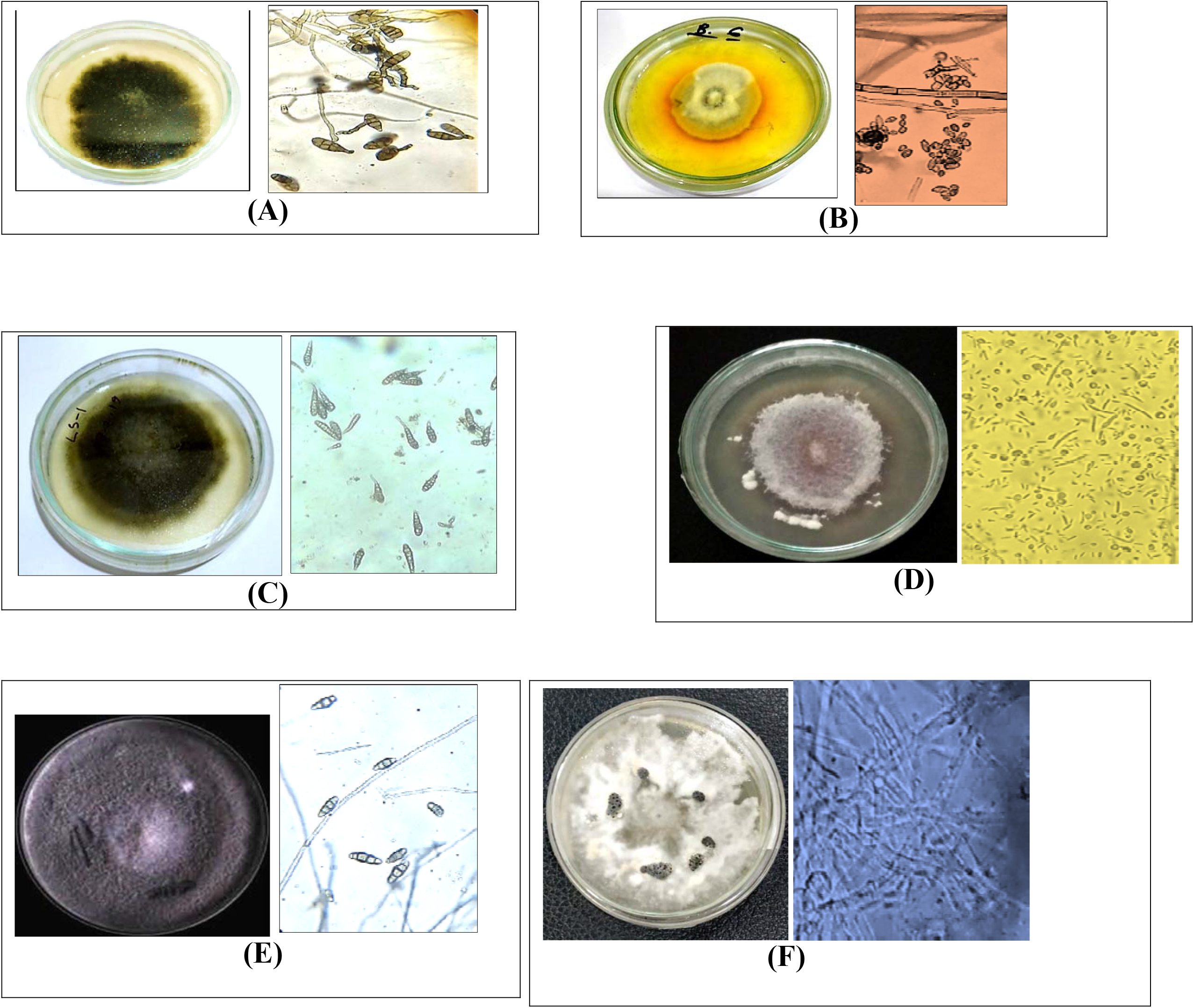
Pure Culture (left) and compound microscopic view 10X (right) of the isolated pathogens. **A**. *Alternaria alternate* isolated from leaf spot disease, **B**. *Botrytis cinerea* isolated from botrytis blight disease, **C**. *Alternaria dianthi* isolated from flower bud rot disease, **D**. *Fusarium oxysporum* isolated from stem rot disease, **E**. *Curvularia lunata* isolated from foliage blight disease, **F**. *Sclerotinia sclerotiorum* isolated from white mold disease

The pathogen responsible for Alternaria leaf spot disease in marigold was identified as *Alternaria* spp. In the case of *Alternaria alternata*, its mycelium was septate, branched, and hyaline when young. Conidiophore was simple, septate, and short, colored, producing conidia at its apex. The dark-colored conidia were both short and long beaked, multicellular, containing several longitudinal and transverse septa, usually coming solitary or in short chains at the tip of the conidiophores. Transverse septa were normally 4-8 with a few longitudinals in each conidium. The shape varied between elliptical, obclavate, or ovoid, tapering to an apex at the distal end. The pure culture of *Alternaria* sp. which was moderately slow-growing, with a colony turning blackish within 10 days on PDA medium.

The pathogen was identified as *Botrytis cinerea*, which causes the Botrytis blight disease. There were characteristic fuzzy gray masses of spores with threadlike, branched hyphal structures, with smooth, brown, tree-like conidiophores about 2 mm long. Numerous globose conidia were present, hyaline and non-septate. A pure culture of *Botrytis* was obtained. The fungus is of medium growth; on PDA culture medium, a whitish, cottony colony was developed within 8 days.

The casual organism was identified as *Alternaria dianthi*, in which the mycelium, when young, was septate, branched, and hyaline. The conidia were dark, with long beaks, and were multicellular with muriform structure, having longitudinal as well as transverse septa. They were usually terminal but sometimes in short chains at the apex of the conidiophores. Conidia had 5-9 transverse septa and several longitudinal septa, were of shape from ellipsoidal to obclavate or ovoid, tapering to a pointed distal end, with the beak often slightly swollen at the tip. A pure culture of *Alternaria dianthi* was obtained from conidia taken from lesions, and colonies on PDA medium were moderately slow-growing in culture and formed a brown to blackish coloration within 10 days.

*Fusarium oxysporum* was isolated as the causal organism of marigold stem rot disease. Under microscope, small, oval-shaped, bicellular microconidia and hyaline, multicellular macroconidia with three septa were seen; the latter were sickle-shaped, notched at the base and at one end. A pure culture of *Fusarium* was isolated and the growth in culture was moderately fast, giving a cottony white appearance on PDA medium within 7 days of incubation.

The casual organisms responsible for foliage blight (flower and leaf blight) in marigold were *Alternaria* sp., *Curvularia lunata*, and *Aspergillus fumigatus*. Colonies of *C. lunata* showed fast growth and were olivaceous to greyish. The conidiophores were darkly pigmented, geniculate with sympodial development. Conidia were curved; sometimes hardly so and arose from an enlarged central cell which was darker than the remaining cells. Large, upright stromata were easily seen in culture. Usually, conidia of *C. lunata* contained three septa and four cells, had an obclavate to elliptical or ovoid appearance, often pointed at the distal end. Colonies of *Curvularia* grew moderately fast in pure culture on PDA medium with a dark white to blackish appearance within 7 days. *Aspergillus fumigatus*: The colonies growing on the PDA plates showed gray-green, cottony colonies on the surface and an off-white reverse. Conidiophores were aseptate, smooth, and greenish with flask-shaped vesicles that are usually fertile over the upper half. Sterigmata were crowded in one series. Conidia were gray-green, one-celled, globose, echinulate and catenate. In *Alternaria* sp. the mycelium is septate, branched, hyaline when young. The conidiophores are simple, septate, short, colored, bearing the conidium at the tip. Conidia dark-colored, beakless to short and long beak-bearing, multicellular, muriform with longitudinal and transverse septa, singly or in short chain at the tips of conidiophores. These conidia contained 4-8 transverse septa and a few longitudinal septa. Their shape ranged from elliptical to obclavate or ovoid, pointed at the distal end. A pure culture of *Alternaria* sp. was established in which, in culture, the colonies grew moderately slowly and showed blackish appearance on PDA medium within 10 days.

White mold disease of marigold is caused by the causal organism *Sclerotinia sclerotiorum*. On pure culture on PDA medium, dark sclerotia were produced. Fragments of dead plants, when kept in a moist chamber, developed white mycelia and sclerotia on them, typical of *S. sclerotiorum*. The fungus produced sclerotia as its survival structure either on or inside the host plant tissues. Since it was documented that *Sclerotium sclerotiorum* had been infective to nearly all forms of plant tissue, ranging from stems to foliage, it also extended to flowers.

### Disease incidence and severity

The tables show that the incidence and severity of diseases varied among locations. Leaf spot was the highest in the Patuapara location at 54.33%, whereas in Hariya, it was relatively low at 30%. Its severity ranged from 9.33 to 18%, out of which Shiorda had the highest. Botrytis blight showed 38.33% incidence in the Sayedpara location and 15.33% in Kuliya, while the severity ranged from 6.67% to 21%. The incidence of flower bud rot was maximum at Mathuapara amounting to 40.66% and minimum in Hariya amounting to 12%, while the severity was minimum in Sadirali at 6.03% and maximum at Mathuapara at 18%. Maximum stem rot infection of 29% was recorded in Nirbashkhola, whereas Shiorda showed the lowest incidence of 7%, while the severity ranged from 2.67 to 13%. The incidence of foliage blight was maximum in Godkhali, 42%, and minimum in Gaburapur, 5.67%, while the severity ranged between 1% and 18%. White mold confined its attack to Jhikargacha, and incidence peaked to a maximum of 26.67% in Shiorda, while it was zero in Patuapara. Severity also varied along the same trend from 0% to 10.67%.

## 4. Discussion

Diseases in marigold were identified from the Jashore district, which is one of the most prevalent districts for marigold in Bangladesh. The field data were collected from natural environments and then analyzed using STATISTIX-10 software. In a thorough survey of the marigold fields all over Bangladesh, six diseases were identified. Of the six diseases identified in this survey, four were generally found at almost every surveyed location, whereas the remaining two were confined to only a few specific areas. The isolated fungi from the diseased plants were *Alternaria alternata, Botrytis cinerea, Alternaria dianthi, Fusarium oxysporum, Curvularia lunata*, and *Sclerotinia sclerotiorum*. These pathogens, reported by previous works, have involvement in marigold diseases. In various growing regions, disease incidence varied from 0% to 54.33%, whereas disease severity was between 1% and 24%. These diseases have also been reported by many scientists, researchers, and plant pathologists around the world.

Alternaria leaf spot, caused by various *Alternaria* species, is a significant disease in marigold, particularly affecting *Tagetes erecta* and *T. patula*. Shamsi and Aktar [23] identified the most virulent pathogen that causes both leaf spot and flower blight of Bangladesh as *Alternaria tagetica*. Similarly, Qui et al. [24] identified *A. tagetica*, which produces phytotoxic compounds such as alternaric acid, that further impairs plant health. Tomioka et al. [25] described the leaf spot and blight, as a newly emerging disease in the region, of marigold caused by necrotic foliar lesions in African and French marigolds. In India, Sen [26] reported *A. zinniae* as a serious threat to African marigold, inducing up to 60% disease severity. Earlier on, Mukerji and Bhasin [27] attributed leaf spot in marigolds to *A. alternata*, while Cotty and Mishaghi [28] observed that *A. tagetica* had already become a serious constraint to high-yielding scented marigold varieties due to the premature defoliation it caused and killing of the plants. This fungus was also found causing inflorescence blight, large tan-to-brown blotches and zonal lesions on leaves. The prevailing environmental factors like pro-longed high moisture, as observed by Hotchkiss and Baxter [29], are highly contributing to aggravating the disease severity caused by *A. tagetica*. Allen et al., [30] observed that Alternaria diseases alone causing defoliation which is more serious and causing maximum loss to vegetative growth, while *A. helianthi* infected maximum at 25° to 28°C with continued presence of water. Shome and Mustafee, [31] reported *A. tagetica*, causing yield loss up to 50-60 percent flowers in marigold and its prevalence was more serious in northern Madhya Pradesh, India.

Botrytis blight is a very destructive disease in *Tagetes erecta* caused by *Botrytis cinerea*. Sultana and Shamsi [32] reported that the crop loss due to this disease is up to 70%. It grows under cool humid conditions and appears as dead blotches on leaves, flowers, and stems. The symptoms include stem rot and collapse of the plants, unopened flower buds, or if opened, the flower decays and falls prematurely. Infected tissues are completely covered by a gray, fuzzy fungal growth with spores. Dhilon and Arora [33] reported that the removal of plant debris, foliage dryness, and avoidance of overhead irrigation were among the control measures against this disease. Disease-resistant cultivars and fungicides would further minimize its infection and spread.

Flower bud rot caused by Alternaria dianthi is a serious disease that affects flower buds and leaves of the plant [34]. Infection is characterized by brown necrotic spots on margins and tips of older leaves, showing advancement into perfect leaf blight. The disease causes young flower buds to shrivel, turn deep brown, and dry up, while mature buds remain slightly green and also fail to open because of the pathogen. Spraying 0.2% Mancozeb and 0.2% Dithane M-45 has been found effective in controlling infection of flower bud rot, showing timely treatment is very important in managing the disease [35].

This highly infectious nature of *Fusarium oxysporum* and *Phytophthora cryptogea* respectively is what causes the severe collar regions of marigold plants, usually leading to the death of the seedlings [34]. As such, at later stages, the disease appears as black streaks along the vascular tissues in the wilting and heavy decaying of roots. During wet weather conditions, salmon-colored spore masses may appear on the infected stems, further aggravating the problem. French marigold and dwarf types are somewhat resistant to the disease, but African marigold types are more susceptible. Tomioka et al. [25] described similar necrotic lesions on *Tagetes erecta* and *T. patula* in Japan where wilting was most noticeable on the orange-flowering varieties. Affected plants had black streaks running upwards from the soil line and pinkish spore masses at the crown which gave evidence of the advanced stage of infection.

An extended study (Shamsi and Aktar [23] isolated a total of 20 fungal species from two marigold species, namely *Tagetes erecta* and *Tagetes patula*. Among these 20 fungal isolates, three major species infecting foliage leaf blight in both the above-mentioned marigold species were identified as *Aspergillus fumigatus, Alternaria alternata*, and *Curvularia lunata*. It generally causes necrotic lesions on leaves, heavy defoliation, and reduces photosynthetic activities, hence weakening plants. Identification of these pathogens gives an indication of their role in the dissemination of the leaf blight and stresses the need for methods to control the disease for a minimum loss of crops and optimum health of plants during marigold cultivation.

Rahman et al. [36] reported white mold disease of marigold caused by *Sclerotinia sclerotiorum*, but it was considered minor. The fungus produces white mold that eventually darkens; infected plant parts that may turn dark green usually appear watery or greasy. Reports in January 2011 showed that the flowers of marigold from Rangpur of Bangladesh had rotten and covered with fluffy white mycelia. Symptoms first appeared on the petals and then progressed to the rest of the flower and the lower portions of the plant. The dark brown lesions containing necrotic tissue were present on the infected leaves and stems, while highly infected plants showed flower drop and wilting of branches.

The present study deals with diversity and prevalence of diseases in marigold plants of Jashore district in Bangladesh. The six predominant diseases of marigold are of fungal origin and were found to have a prevalence ranging from 0% to 54.33% through in-depth analysis. Leaf spot, botrytis blight, foliage blight, and flower bud rot are major diseases that seriously threaten this crop. This study identified specific pathogens responsible for diseases, hence the reason targeted control strategies are highly essential. Apparent spatial differences in disease occurrence across the tested regions also signal region-specific management practices and support the application of IPM strategies for sustainable disease management of marigold cultivation. In summary, the present study has furthered our understanding of the prevalence and impact of the field diseases affecting marigold cultivation in the Jashore district of Bangladesh. Identification of the major disease problems and their respective pathogens provides the necessary information to guide development of efficient disease management practices, an integral component of sustainable marigold flower production in the area.

## 5. Conclusions

This experiment utilized cultural and morphological methods for detecting six different diseases affecting marigold, studying their prevalence and severity in different places. The results showed a wide variation in the prevalence and severity of the diseases, with leaf spot, Botrytis blight, foliage blight (including flower and leaf blight), and flower bud rot of major importance to marigold cultivation in the Jashore district of Bangladesh. Overcoming these will require adequate management of the diseases to reduce the potential losses of crops. This is significant research, though it also points out the need to continue in-depth analysis of the underlying conditions that drive the dynamics of diseases, such as climate change and agricultural practices. Subsequent studies should target such aspects in order to come up with more specific management strategies. Resistant varieties of marigold will also be discovered, and monitoring of diseases over a long period is highly essential for the success of sustainable agriculture. This study thus forms a very important prelude to further studies on disease management strategies to be conducted toward better sustainability and resilience in marigold cultivation in Bangladesh.

## Ethical approval

This study does not include human or animal subjects.

## Data Availability Statement

All relevant data are available in the study.

## Acknowledgments

The author would like to express gratitude to the Ministry of Science and Technology, Government of Bangladesh, for awarding the National Science and Technology (NST) Fellowship that supported this research in the 2018-19 session. She is also grateful to the Supervisor for providing research materials and facilities through the Grant for Advance Research in Education (GARE) sponsored by the Ministry of Education, Government of the People’s Republic of Bangladesh. This support was given under the project entitled “Establishment of Plant Disease Clinic at Sher-e-Bangla Agricultural University, Dhaka, Bangladesh.”

## Author contribution statement

AS = Conceptualization, Investigation, Writing - original draft, Data curation, Software, Methodology, Funding Acquisition, Project Administration, Formal analysis, Writing - review & editing, Visualization, Resources, Validation. ANFA = Supervision, Conceptualization, Methodology, Writing – Original Draft, Data curation, Validation; RF = Methodology, Resources, Visualization, Writing - review & editing, Formal analysis, Validation. All authors have reviewed the manuscript and provided their feedback.

